## Supplementary Information for "Structure of human MUTYH and functional profiling of cancer-associated variants reveal an allosteric network between its [4Fe-4S] cluster cofactor and active site required for DNA repair"

**Table of Contents**

**Supplementary Methods:** design of MBP-MUYTH protein for crystallography.

**Figure S1.-** Details of the pET28-MBP-MUTYH construct, MUTYH protein purification and characterization.

**Figure S2.-** Multiple sequence alignment of MutY/MUTYH amino acid sequences.

**Figure S3.-** Structural comparison of mMutyh/MUTYH.

**Figure S4**. Structural mapping of cancer-associated variants (CAVs) onto the MUTYH-TSAC structure.

**Figure S5.-** Representative results of the qualitative glycosylase characterization of MBP-MUTYH cancer-associated mutants.

**Figure S6.** Representative qualitative binding assays.

**Figure S7.-** Structural analysis of the alternative conformations of the [4Fe-4S] cluster within the R149Q GsMutY-THF:OG structures.

**Table S8.-** Adenine Glycosylase Actvity of WT GsMutY and R149Q GsMutY.

**Figure S9.-** Conservation of structural interplay between the [4Fe-4S] cluster and the active site in Helix-hairpin-Helix DNA glycosylases.

**Table S10**. Adenine Glycosylase Activity, Substrate and Product Analog Affinity, Metal analysis of cancer-associated mutants within the [4Fe-4S] cluster of MUTYH.

**Table S11.** Mutation suppression activity of cancer-associated mutants within the [4Fe-4S] cluster of MUTYH.

**Table S12.** Reduction of restraints throughout equilibration during molecular dynamic simulations.

**Figure S13.-** Results of the molecular dynamic simulations of WT MUTYH, N238S and R241Q cancer-associated mutants in human structure.

**Figure S14.-** Energy decomposition analysis of the nonbonded interactions of the MUTYH human structure with respect to the [4Fe-4S] cluster.

**Figure S15.-** Results of the molecular dynamic simulations of WT, N209S and R212Q cancer-associated mutants in mMutyh mouse structure.

**Figure S16.-** Results of the molecular dynamic simulations of WT, N238S and R241Q cancer-associated mutants in mouse Mutyh structure.

**Table S17.-** Data collection and model refinement statistics for MUTYH-TSAC and R149Q Gs MutY structures.

**Zip file contains Normal Mode Analysis videos and Multiple Sequence Alignment used for MutY/MUTYH coevolutionary analysis.**

**Supplementary Methods: Design of MBP-MUYTH protein for crystallography.**

Open Reading Frame encoding the fusion MBP-MUYTH protein. Highlighted in red and green are the region encoding the Maltose binding protein region and the codon-optimized *MUTYH* gene optimized for its expression in *E. coli*, respectively. A linker with three tandem repeats of Gly-Ser is shown in pink followed by TEV cleavage site in blue. NcoI, NdeI and NotI restrictions sites used for cloning are underlined.

**CCATGG**GCAAAATCGAAGAAGGTAAACTGGTAATCTGGATTAACGGCGATAAAGGCTATAACGGTCTCGCTGAAGTCGGTAAGAAATTCGAGAAAGATACCGGAATTAAAGTCACCGTTGAGCATCCGGATAAACTGGAAGAGAAATTCCCACAGGTTGCGGCAACTGGCGATGGCCCTGACATTATCTTCTGGGCACACGACCGCTTTGGTGGCTACGCTCAATCTGGCCTGTTGGCTGAAATCACCCCGGACAAAGCGTTCCAGGACAAGCTGTATCCGTTTACCTGGGATGCCGTACGTTACAACGGCAAGCTGATTGCTTACCCGATCGCTGTTGAAGCGTTATCGCTGATTTATAACAAAGATCTGCTGCCGAACCCGCCAAAAACCTGGGAAGAGATCCCGGCGCTGGATAAAGAACTGAAAGCGAAAGGTAAGAGCGCGCTGATGTTCAACCTGCAAGAACCGTACTTCACCTGGCCGCTGATTGCTGCTGACGGGGGTTATGCGTTCAAGTATGAAAACGGCAAGTACGACATTAAAGACGTGGGCGTGGATAACGCTGGCGCGAAAGCGGGTCTGACCTTCCTGGTTGACCTGATTAAAAACAAACACATGAATGCAGACACCGATTACTCCATCGCAGAAGCTGCCTTTAATAAAGGCGAAACAGCGATGACCATCAACGGCCCGTGGGCATGGTCCAACATCGACACCAGCAAAGTGAATTATGGTGTAACGGTACTGCCGACCTTCAAGGGTCAACCATCCAAACCGTTCGTTGGCGTGCTGAGCGCAGGTATTAACGCCGCCAGTCCGAACAAAGAGCTGGCAAAAGAGTTCCTCGAAAACTATCTGCTGACTGATGAAGGTCTGGAAGCGGTTAATAAAGACAAACCGCTGGGTGCCGTAGCGCTGAAGTCTTACGAGGAAGAGTTGGTGAAAGATCCGCGTATTGCCGCCACTATGGAAAACGCCCAGAAAGGTGAAATCATGCCGAACATCCCGCAGATGTCCGCTTTCTGGTATGCCGTGCGTACTGCGGTGATCAACGCCGCCAGCGGTCGTCAGACTGTCGATGAAGCCCTGAAAGACGCGCAGACTAATTCGGGATCTGGCAGTGGTTCTGAGAATCTTTATTTTCAGGGC**CATATG**CAGGCTGCCTCACAGGAAGGCCGCCAGAAACACGCAAAGAATAACAGTCAGGCAAAACCAAGCGCTTGCGATGGCTTGGCGCGTCAACCGGAGGAAGTCGTGTTACAAGCTAGTGTCTCTTCATATCACCTTTTCCGTGACGTTGCAGAAGTTACAGCTTTCCGCGGTTCTTTATTAAGTTGGTATGACCAGGAAAAGCGCGACTTACCTTGGCGTCGCCGCGCGGAGGATGAAATGGACCTTGACCGTCGTGCTTATGCCGTGTGGGTATCGGAGGTTATGTTGCAACAGACGCAGGTAGCCACCGTGATTAACTACTACACGGGTTGGATGCAAAAGTGGCCAACGTTGCAGGATCTGGCGTCCGCTTCACTGGAAGAAGTGAATCAGTTATGGGCAGGATTAGGGTACTACTCGCGCGGTCGTCGTCTGCAAGAGGGGGCACGCAAAGTTGTTGAAGAATTGGGAGGCCACATGCCGCGCACTGCAGAAACCTTACAACAGTTACTTCCCGGGGTAGGCCGTTATACCGCTGGAGCAATTGCATCCATTGCTTTCGGACAGGCTACTGGAGTAGTTGATGGAAATGTGGCACGTGTTTTGTGTCGCGTTCGCGCCATCGGCGCAGACCCTTCATCCACTTTGGTATCTCAGCAGCTGTGGGGATTAGCACAGCAGTTAGTCGATCCCGCTCGTCCCGGCGACTTCAACCAAGCGGCCATGGAACTTGGTGCCACGGTATGTACGCCACAGCGTCCGCTTTGTAGCCAGTGTCCGGTTGAGAGCTTGTGTCGTGCTCGCCAACGTGTAGAGCAGGAACAACTTTTAGCAAGCGGTTCCCTGTCTGGCTCTCCCGATGTTGAAGAATGTGCACCTAACACAGGTCAGTGTCATCTTTGTCTTCCGCCGAGTGAACCCTGGGATCAGACGTTGGGCGTCGTGAACTTTCCGCGTAAGGCCTCACGCAAACCCCCTCGCGAGGAAAGTAGTGCAACGTGCGTACTGGAGCAACCAGGAGCTTTAGGCGCGCAAATTTTGCTTGTTCAACGCCCGAATAGTGGTCTGTTAGCTGGCCTGTGGGAATTTCCATCAGTAACGTGGGAGCCCAGCGAGCAGCTGCAACGCAAAGCCTTATTGCAAGAGTTACAACGCTGGGCTGGACCGTTGCCAGCGACTCATCTGCGTCATTTAGGAGAAGTAGTACATACGTTCAGTCACATCAAGCTGACCTATCAAGTTTATGGCCTTGCCTTAGAAGGGCAAACTCCGGTGACCACCGTGCCGCCCGGCGCACGTTGGCTTACACAAGAAGAATTCCACACGGCTGCTGTCAGTACGGCGATGAAGAAGGTCTTTCGCGTGTATCAGGGGCAGCAACCTGGAACGTGTATGGGTTCAAAACGTAGTCAAGTATCGAGTCCATGTTCTCGTAAGAAGCCACGTATGGGTCAGCAGGTCCTGGATAACTTTTTCCGTAGTCATATCTCCACAGACGCCCATAGCTTGAACTCGGCGGCCCAGTCAGGCGAGAATCTTTATTTTCAGGGCAGCGGATCAGGGAGCGGATCAGGCAGCGGGCATCACCATCACCATCACCATCACTGA**GCGGCCGC**

**Figure S1.-** Multiple sequence alignment of MutY/MUTYH amino acid sequences. The protein sequences included in the analysis were *Escherichia coli* MutY (EcMutY; NCBI ID ABV07358.1), *Geobacillus stearothermophilus* MutY (GsMutY; WP_013522988.1), human MUTYH (NP_001121897.1) and mouse Mutyh (NP_001153053.1). Catalytic residues are colored in yellow, positions studied herein in green, FSH Loop implied in OG recognition in gray, and IDC region in red. Residues involved in the structural interplay between the [4Fe-4S] cluster and active site are indicated with asterisks. Residues within the Zinc linchpin motif are highlighted in bold letters.

```
          1        10        20        30        40        50        60
          |        |         |         |         |         |         |
 EcMutY   ------------------------------------------------------------
 GsMutY   ------------------------------------------------------------
  MUTYH   MTPLVSRLSRLWAIMRKPRAAVGSGHRKQAASQEGRQKHAKNNSQAKPSACDACAGMIAE
  Mutyh   --------------MKKLQASVRS-HKKQPANHKRRRTRALSSSQAKPSSLD--------

 EcMutY   ------------------------------MQASQFSAQVLDWYDKYGRKTLPWQI----
 GsMutY   -----------------------MTRETERFPAREFQRDLLDWFAR-ERRDLPWRK----
  MUTYH   CPGAPAGLARQPEEVVLQASVSSYHLFRDVAEVTAFRGSLLSWYDQ-EKRDLPWRRRAED
  Mutyh   ------GLAKQKREELLQASVSPYHLFSDVADVTAFRSNLLSWYDQ-EKRDLPWRNLAKE

 EcMutY   ----DKTPYKVWLSEVMLQQTQVATVIPYFERFMARFPTVTDLANAPLDEVLHLWTGLGY
 GsMutY   ----DRDPYKVWVSEVMLQQTRVETVIPYFEQFIDRFPTLEALADADEDEVLKAWEGLGY
  MUTYH   EMDLDRRAYAVWVSEVMLQQTQVATVINYYTGWMQKWPTLQDLASASLEEVNQLWAGLGY
  Mutyh   EANSDRRAYAVWVSEVMLQQTQVATVIDYYTRWMQKWPKLQDLASASLEEVNQLWSGLGY

                                                                  * *
 EcMutY   YARARNLHKAAQQVATLHGGKFPETFEEVAA-LPGVGRSTAGAILSLSLGKHFPILDGNV
 GsMutY   YSRVRNLHAAVKEVKTRYGGKVPDDPDEFSR-LKGVGPYTVGAVLSLAYGVPEPAVDGNV
  MUTYH   YSRGRRLQEGARKVVEELGGHMPRTAETLQQLLPGVGRYTAGAIASIAFGQATGVVDGNV
  Mutyh   YSRGRRLQEGARKVVEELGGHMPRTAETLQQLLPGVGRYTAGAIASIAFDQVTGVVDGNV

           *                                                *
 EcMutY   KRVLARCYAVSGWPGKKEVENKLWSLSEQVTPAVGVERFNQAMMDLGAMICTRSKPKCSL
 GsMutY   MRVLSRLFLVTDDIAKPSTRKRFEQIVREIMAYENPGAFNEALIELGALVCTPRRPSCLL
  MUTYH   ARVLCRVRAIGADPSSTLVSQQLWGLAQQLVDPARPGDFNQAAMELGATVCTPQRPLCSQ
  Mutyh   LRVLCRVRAIGADPTSTLVSHHLWNLAQQLVDPARPGDFNQAAMELGATVCTPQRPLCSH

 EcMutY   CPLQNGCIA-----------------------------------AANNSW------ALYP
 GsMutY   CPVQAYCQAFAEGVA-----------------------------------------EELP
  MUTYH   CPVESLCRARQRVEQEQLLASGSLSGSPDVEECAPNTGQCHLCLPPSEPWDQTLGVVNFP
  Mutyh   CPVQSLCRAYQRVQRGQL---SALPGRPDIEECALNTRQCQLCLTSSSPWDPSMGVANFP

 EcMutY   GKKPKQTLPE-RTGYFLLLQH----EDEVLLAQRPPSGLWGGLYCFPQFADEES------
 GsMutY   VKMKKTAVKQVPLAVAVLADD----EGRVLIRKRDSTGLLANLWEFPSCETDGADGKEK-
  MUTYH   RKASRKPPREESSATCVLEQPGAL-GAQILLVQRPNSGLLAGLWEFPSVTWEPSEQLQRK
  Mutyh   RKASRRPPREEYSATCVVEQPGAIGGPLVLLVQRPDSGLLAGLWEFPSVTLEPSEQHQHK

 EcMutY   -----LRQWLAQ---RQIAADNLTQ-LTA--FRHTFSHFHLD----------IVPM----
 GsMutY   -----LEQMVGE---QYGLQVELTEPIVS--FEHAFSHLVWQLTVFPGRLVHGGPVEE-P
  MUTYH   ALLQELQRWAGP-----LPATHLRH-LGE--VVHTFSHIKLTYQVYGLALEGQTPVTTVP
  Mutyh   ALLQELQRWCGP-----LPAIRLQH-LGE--VIHIFSHIKLTYQVYSLALD-QAPASTAP

 EcMutY   ----WLPVSSF-----TGCMDEGNALWYNLAQPPSVG-----LAAPVER-----------
 GsMutY   --YRLAPEDELKAYAFPVSHQRVWREYKEWASG---------VRPPP-------------
  MUTYH   PGARWLTQEEFHTAAVSTAMKKVFRVYQGQQPGTCMGSKRSQVSSPCSRKKPRMGQQVLD
  Mutyh   PGARWLTWEEFCNAAVSTAMKKVFRMYEDHRQGTRKGSKRSQVCPPSSRKKPSLGQQVLD

 EcMutY   --LLQQLRTGAPV------
 GsMutY   -------------------
  MUTYH   NFFRSHISTDAHSLNSAAQ
  Mutyh   TFFQRHIPTDKPNSTTQ--
```

**Figure S2.-** Structural comparison of Mutyh/MUTYH. Structural alignment including both copies of MUTYH-DNA complexes included the unit cell and mouse Mutyh (PDB ID; 7EF8)^1^. A close-up view of the catalytic residues, OG recognition sphere and [4Fe-4S] cluster is shown. Moreover, a schematic amino acid sequence analysis is displayed indicating percent identity per regions.

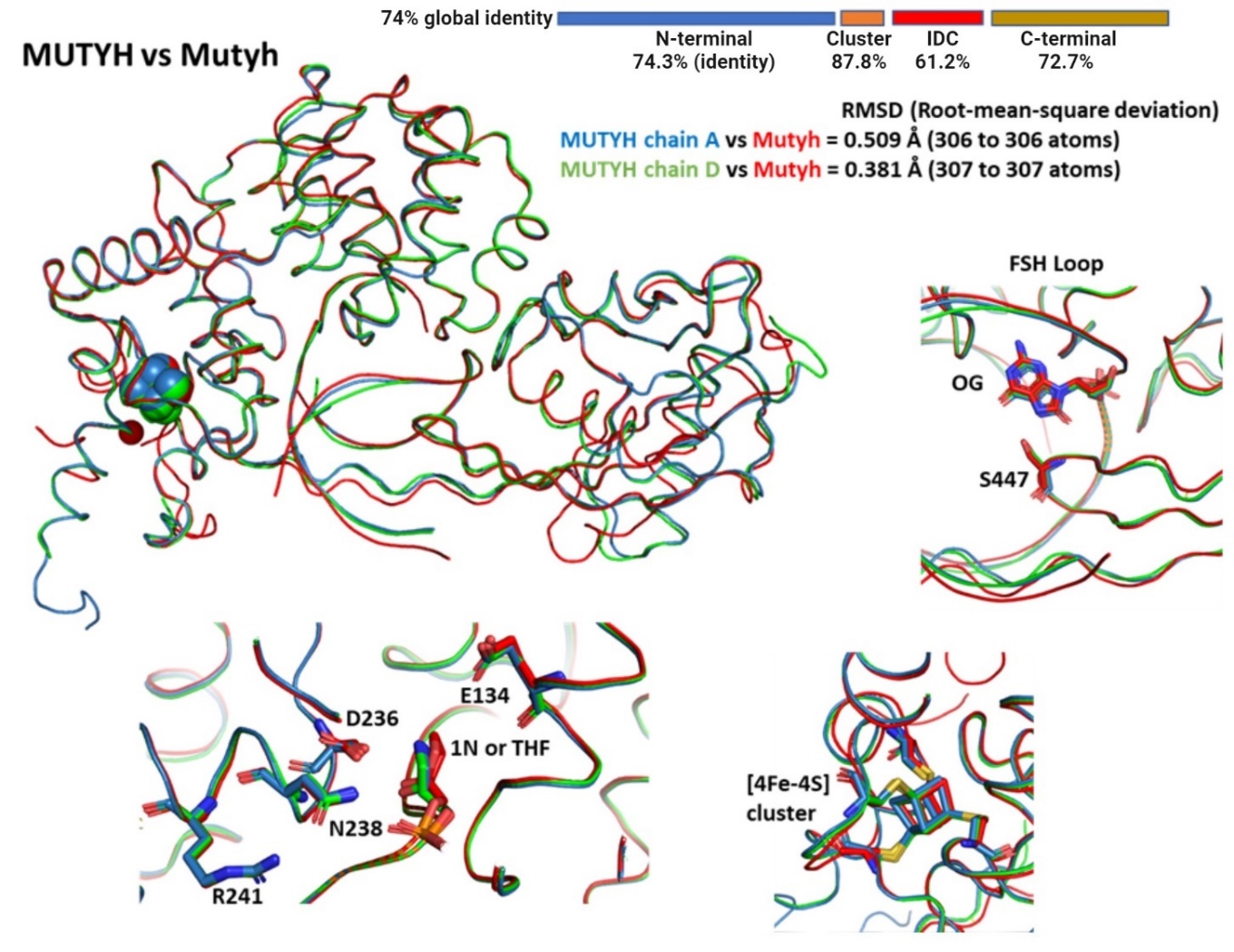

Here, we compared the human MUTYH structure and sequence to the mouse Mutyh to determine if the absence of the Zn ion resulted in overall or localized structural differences. Both structures, in spite of being only 74% identical in amino acid sequence, aligned with a RMSD of up to 0.381 Å. However, in the IDC region the identity drops to 61% (figure S2) and this may impact the structure, flexibility and Zn lability that results in the difference localized here between the MUTYH and mouse Mutyh structures. Importantly, catalytic residues (Asp236 and Glu134), the residue involved in the recognition and stabilization of OG (Ser447), and even the coordination sphere for the [4Fe-4S] cluster and the [4Fe-4S] cluster itself in MUTYH align perfectly with its mouse Mutyh counterpart. Thus, we captured an active conformation of MUTYH which can be reliably used to infer structure-activity relationships.

**Figure S3.-** Details of the pET28-MBP-MUTYH construct, MUTYH protein purification and characterization. Upper-left panel; Scheme of the MBP-MUTYH/pET28 construct used in this work. Upper-right panel; Crystals of MUTYH-TSAC. The MUTYH-DNA complex formation for crystallization was carried with 118 μM MUTYH-DNA (1N:OG) complex. The best crystals in terms of size and X-ray diffraction quality were obtain in 0.1 M Bis-Tris [pH 5.5], 0.2 M Ammonium Sulphate and 20% PEG 3350. Lower-left panel; results of the purification of MUTYH protein. First lane, molecular weight ladder. Second lane, elution fraction after Nickel affinity chromatography. Third lane, Elution fraction after TEV cleavage step. Fourth lane, pure MUTYH after Heparin affinity chromatography. Lower-right panel. Biochemical characterization of MUTYH without tags or fusion MBP protein.

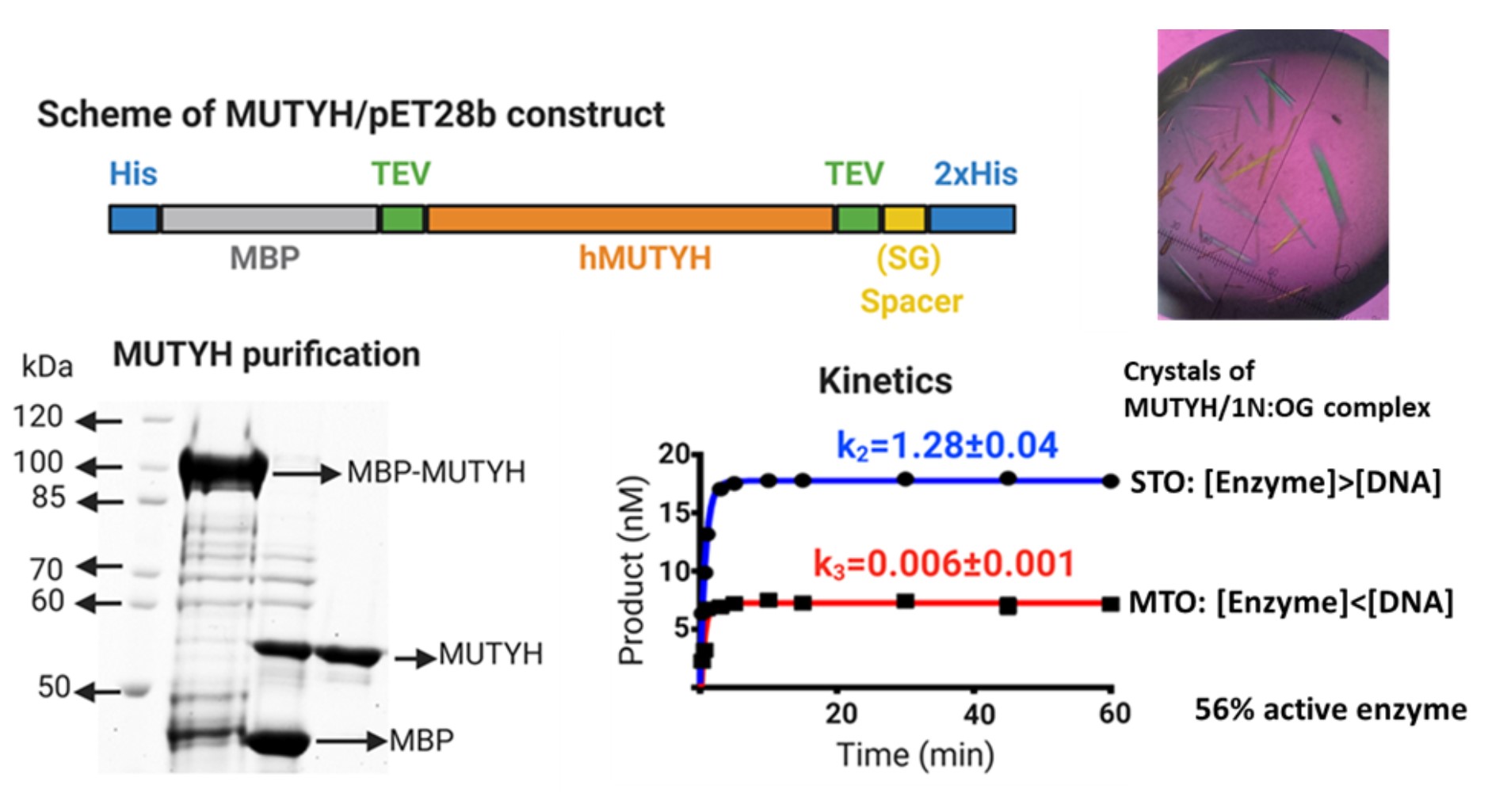

The MUTYH overexpression protocol involved bacterial overexpression in TB at 15 °C, using a BL21 (DE3) strain containing plasmids encoding folding chaperones (pKJE7) and [4Fe-4S] cluster assembly machinery (pKRISC). This method circumvented MUTYH toxicity in *E. coli* and provided yields of >5 mg of MUTYH/L of culture. The crystallization conditions were optimized from those reported for mMutyh^1^ with a key difference being that we used a DNA duplex (11-nucleotide duplex with overhanging ends) containing an azaribose (1N) transition state analog across OG (Figure 1A). The use of 1N, which mimics the oxacarbenium ion transition state/intermediate in terms of shape and charge^1^, provided a highly stable DNA-MUTYH complex that crystallized overnight and diffracted at high resolution (1.9 Å). The solved MUTYH structure showed two copies of DNA-MUTYH complexes per unit cell (table S16).

**Figure S4**. Structural mapping of cancer-associated variants (CAVs) onto the MUTYH-TSAC structure.

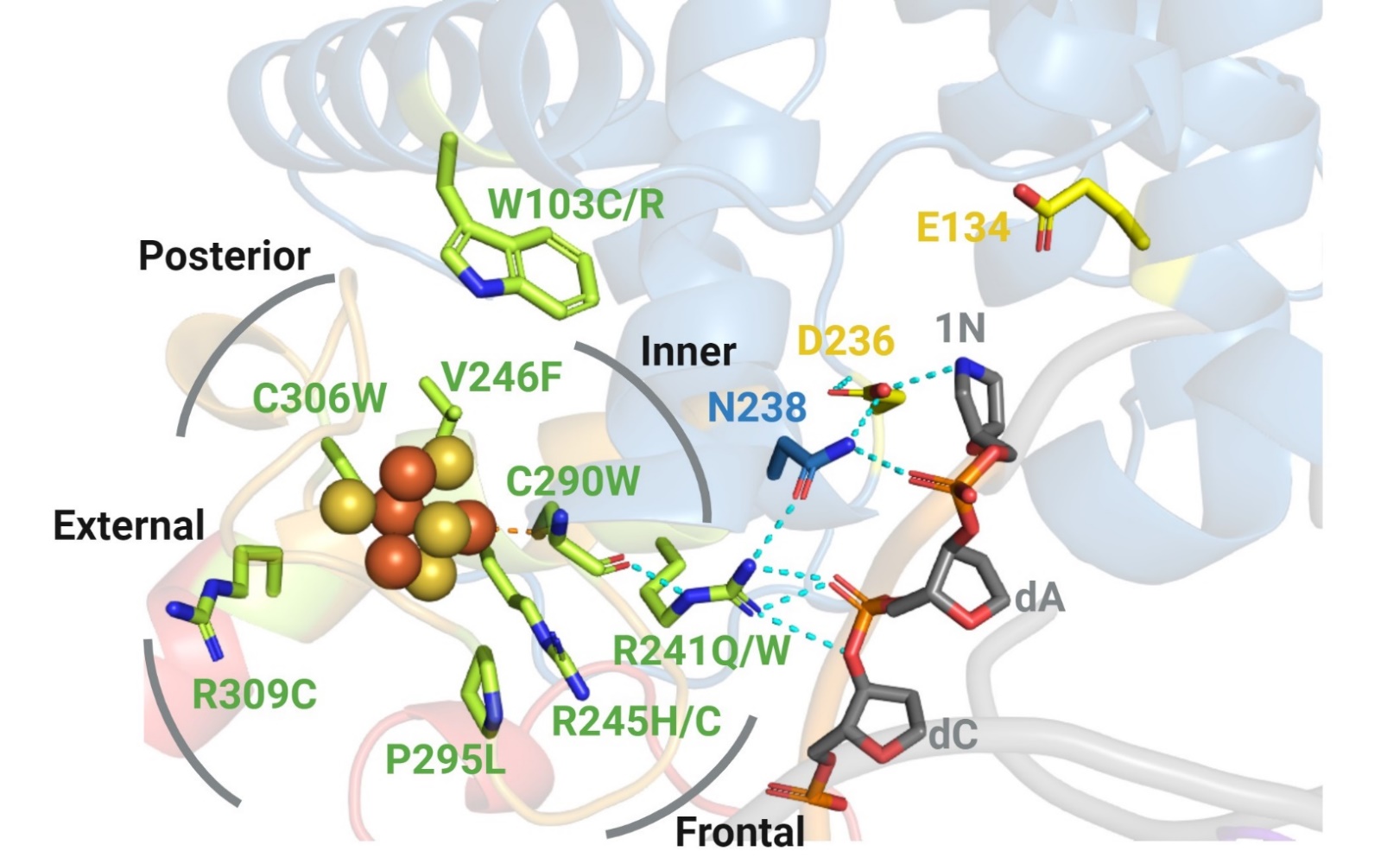

Mapping MAP missense variants onto the MUTYH-TSAC structure was performed to provide insight into the potential consequences of the amino acid alterations. Initially, we identified 10 MAP variants within the [4Fe-4S] cluster binding domain and FCL motif reported in the Leiden Open (source) Variation Database (LOVD 3.0)^2^; W103R, R241Q/W, R245H/C, V246F, C290C, P295L, C306W and R309CW. An additional mutation at Trp103, W103C, was predicted as a pathogenic somatic mutation found in ovarian cancer in the COSMIC database^3^. Cys290 and Cys306 are two of the four cysteines (Cys290, Cys297, Cys300 and Cys306) coordinating the [4Fe-4S] cluster. The nonconservative Cys→Trp mutation of these positions would be anticipated to be destabilizing due to inability to coordinate the cofactor and steric bulk. The Cys290 ligand anchors one end of the solvent exposed to Iron-Sulfur Loop (FCL) motif that interacts with DNA; in addition, Pro295, associated with MAP variant P295L, is also located in the FCL resulting in a structural kink that is likely functionally significant based on the high degree of conservation at this position. The Cys306 ligand is located at posterior face of the [4Fe-4S] cluster packed against several alpha helices in the catalytic domain. CAV V246F and R245H/C are located near Cys306, shielding the cluster from solvent, and providing for hydrophobic packing within the catalytic domain. Notably, the C^α^ of Arg245 is at the posterior face of the [4Fe-4S] cluster but its side chain is projected toward the frontal face. The variant R309C is located on the solvent exposed external lateral side of this metal cofactor, and mapped at the inner lateral side is the R241Q/W MAP variants. Trp103 is located at the interface of the [4Fe-4S] cluster motif and catalytic pocket, and the nonconservative mutations of Trp103 to Arg and Cys, reported as MAP and cancer-associated somatic mutations, respectively, would be anticipated to destabilize this interface, altering both protein stability and catalysis.

**Figure S5.** Representative results of the qualitative glycosylase characterization of MBP-MUTYH cancer-associated mutants. Condition used: 20 nM of A:OG-containing DNA duplex was titrated with increasing concentrations of MBP-MUTYH (10-250 nM) for 1 h at 37 °C. The reactions were resolved by UREA-PAGE and visualized by storage phosphor autoradiography.

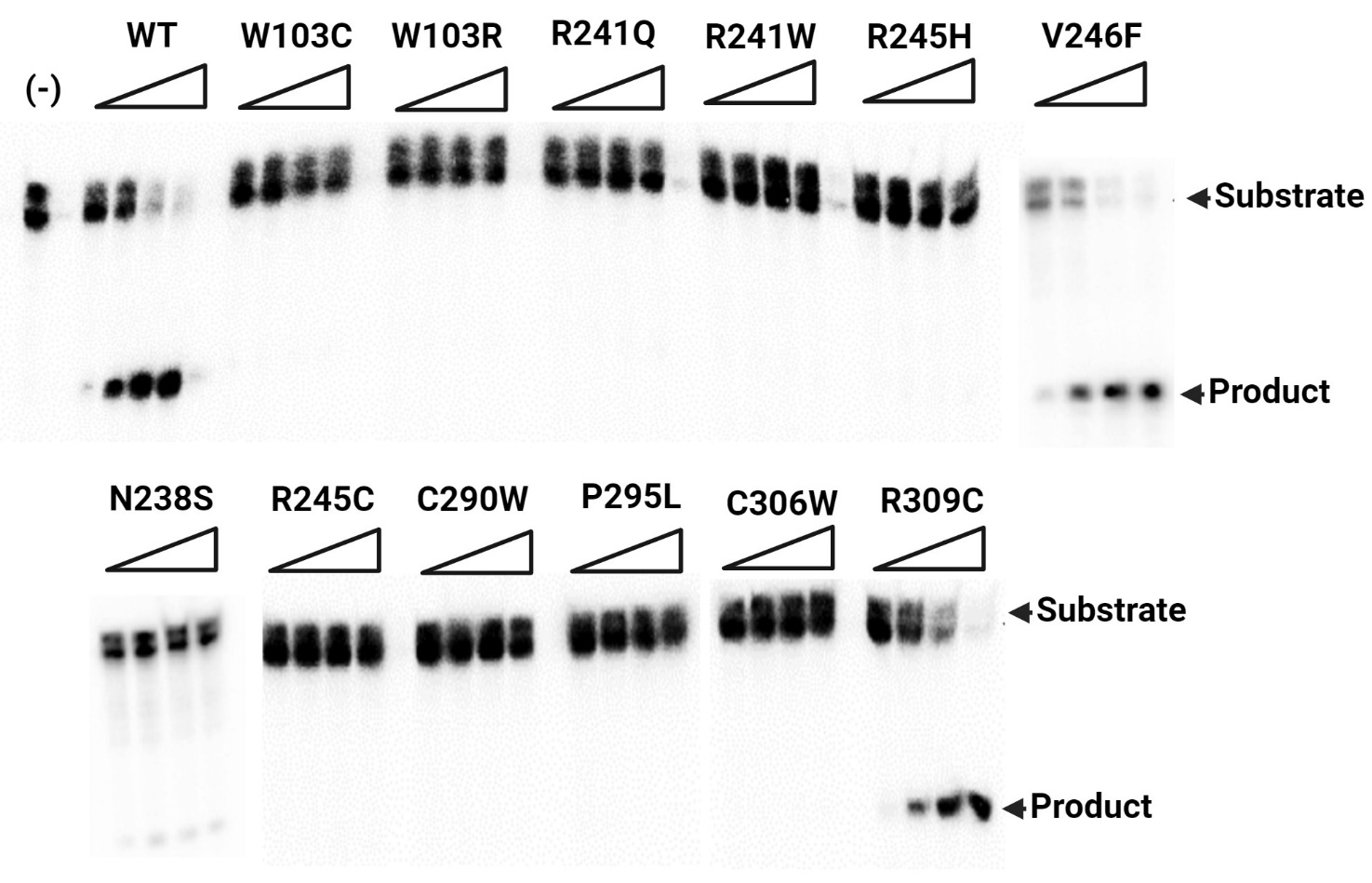

**Figure S6.** Representative qualitative binding assays. 20 nM of THF:OG-containing DNA duplex was titrated with increasing concentrations of MBP-MUTYH (6.25-400 nM) for 20 min at 25 °C. Binding reactions were resolved by native-PAGE

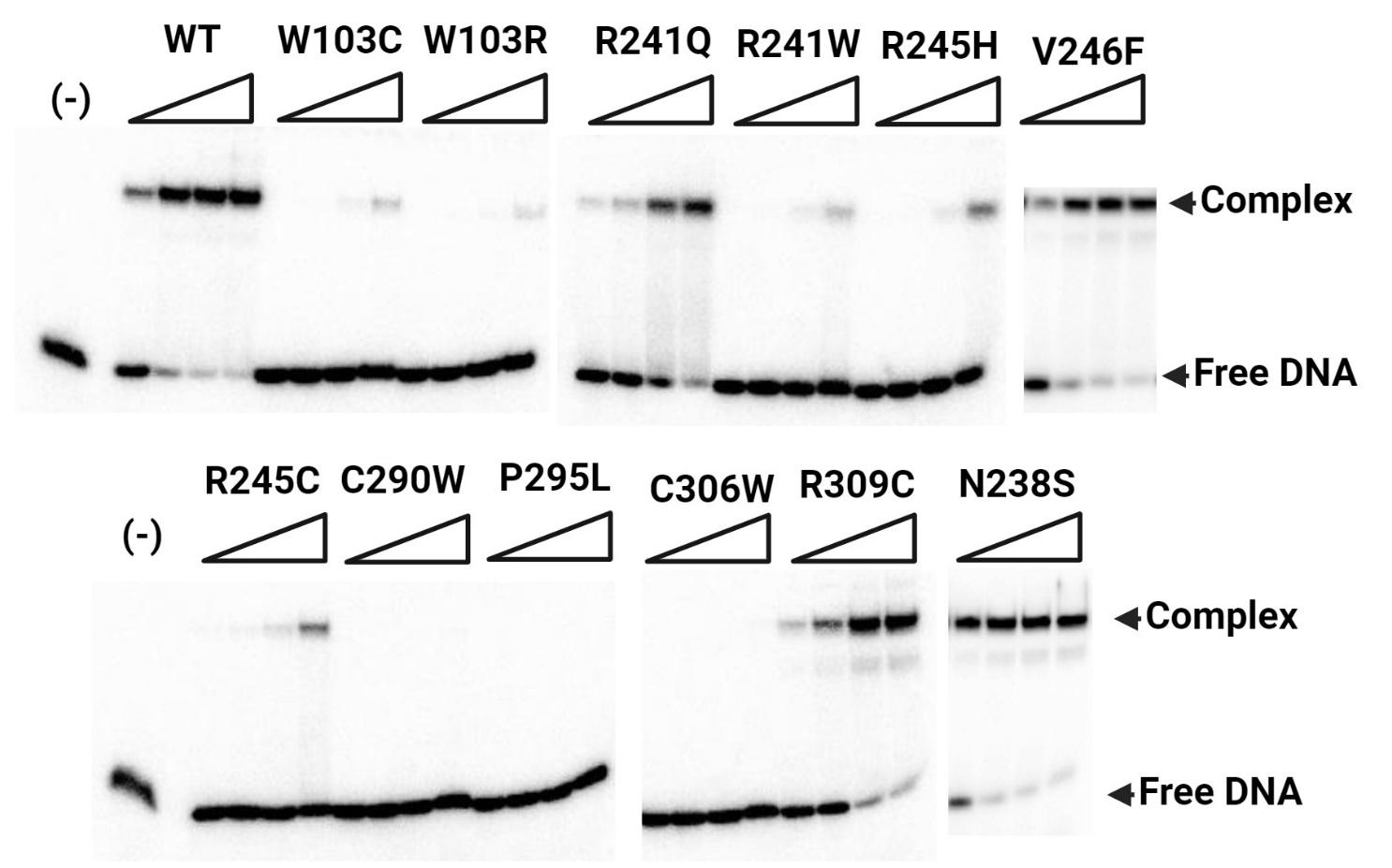

**Figure S7.-** Structural analysis of the alternative conformations of the [4Fe-4S] cluster within the R149Q GsMutY-THF:OG structures. The 2|Fo|–|Fc| omit map (blue) was calculated to the 1.58 Å resolution limits, and contoured at 1.0 rmsd. A) The ANOM map contoured to 5.0 rmsd (gold) shows elongated anomalous signal for the [4Fe-4S] cluster indicating alternative conformations of the cofactor. Difference map of B) [4Fe-4S] cluster motif modeling two conformation of the cofactor and C) after removing the A conformer.

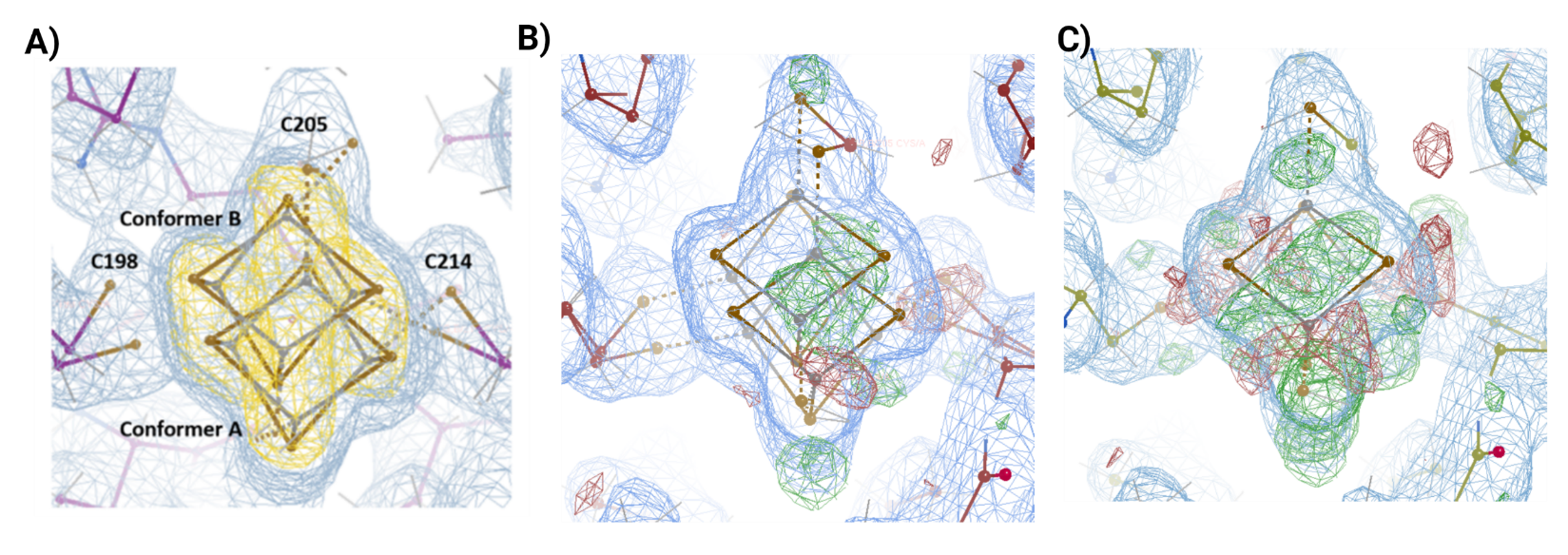

| **Table S8.-** Adenine Glycosylase Activity of WT GsMutY and R149Q GsMutY. | | | | |
| --- | --- | --- | --- | --- |
| Enzyme | Active fraction  (%) | k_2_ (min^-1^)  OG:A | k_2_ (min^-1^)  OG:P | k_3_ (min^-1^)  OG:A |
| WT Gs MutY | 39±1^b^ | 54±4^b^ | 1.13±0.03^c^ | 0.035±0.004^d^ |
| R149Q Gs MutY | 47±5 | 1.11±0.06 | 0.15±0.02 | 0.007±0.001 |
| ^a^The glycosylase assays conducted to measure *k_2_* values were done at 60 °C with 165 nM enzyme, 20 nM radiolabeled DNA, and 30 mM NaCl. The glycosylase assays conducted to measure *k_3_* values were done at 60 °C. ^b,c,d^ Previously reported^4-6^. | | | | |

**Figure S9.-** Conservation of the residues that participate in the allosteric network connecting the [4Fe-4S] cluster and the active site in Helix-hairpin-Helix DNA glycosylases. A close-up view of the residues involved in the structural connectivity is shown. The H-bond network is shown in yellow dotted lines. The structures shown are human MUTYH, MIG (PDB ID 1KEA)^7^ and EndoIII (1ORP)^8^. On the right side a logo sequence of the corresponding region involved in the interplay highlighting the residues that participates in the H-bond network (orange boxes).

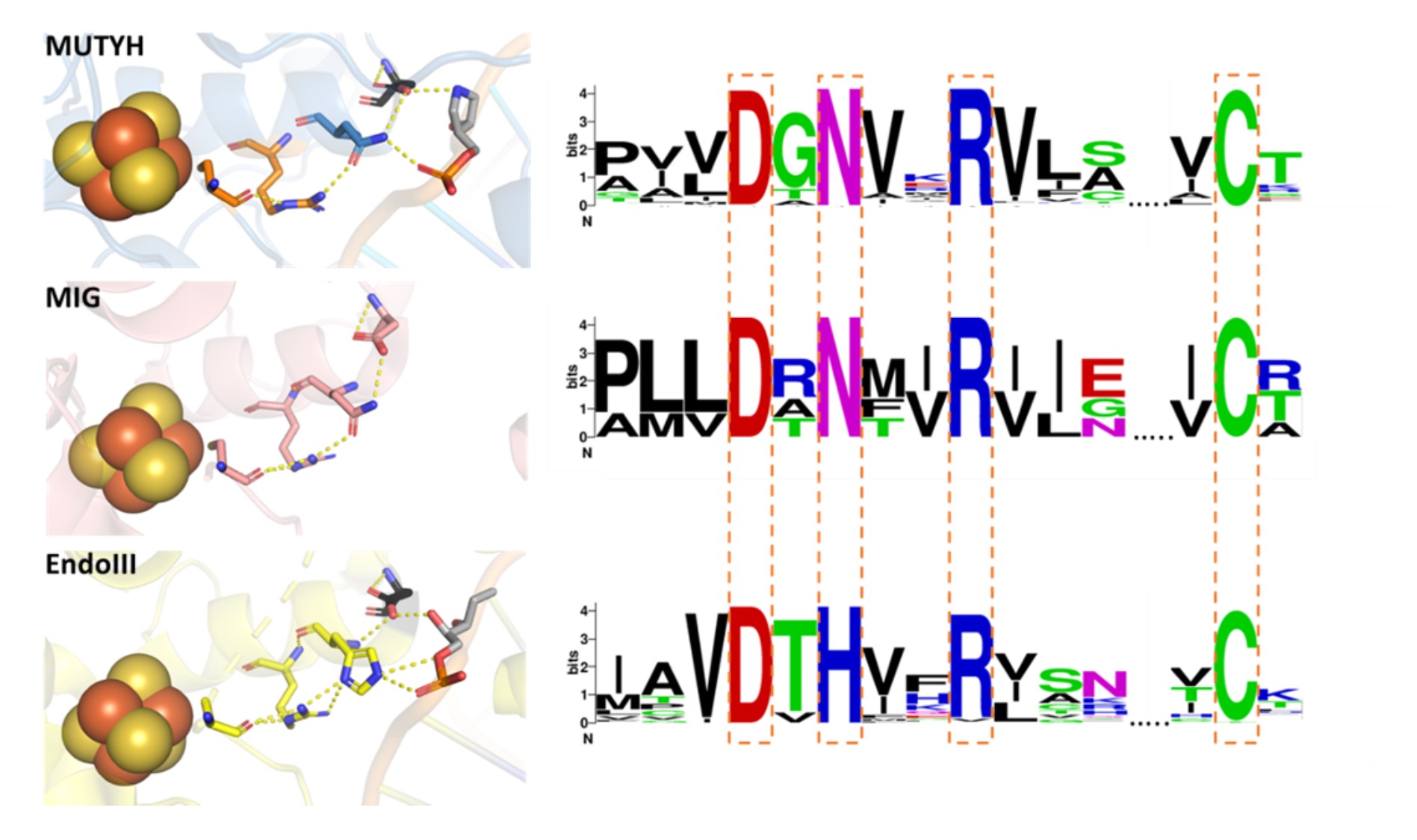

| Table S10. Adenine Glycosylase Activity^a^, Substrate and Product Analog Affinity^a^ ,metal analysis^b^ of cancer-associated mutants within the [4Fe-4S] cluster of MBP-MUTYH. | | | | | | | |
| --- | --- | --- | --- | --- | --- | --- | --- |
| MUTYH | [4Fe-4S] load | Zn load | *k_2_* (min^-1^)  A:OG | *k_3_* (min^-1^)  A:OG | Active MUTYH | *k_D_* (nM)  fA:OG | *k_D_* (pM)  THF:OG |
| WT (MBP free) | 5.1±0.1 | 1.2±0.1 | 1.3±0.1 | 0.006±0.001 | 56-35% | ND | ND |
| WT | 4.4±0.1 | 0.2 ±0.1 | 1.2±0.1 | 0.019±0.008 | 20 % | 32±4 | <10 |
| W103C | 0.0 | 0.0 | ND | ND | ND | ND | ND |
| W103R | 0.0 | 0.0 | ND | ND | ND | ND | ND |
| N238S | 4.1±0.2 | 0.3±0.1 | ND | ND | ND | 124±10 | 46±10 |
| R241Q | 4.4±0.1 | 0.10±0.02 | ND | ND | ND | 629±80 | 467±70 |
| R241W | 0.0 | 0.0 | ND | ND | ND | ND | ND |
| R245H | 0.0 | 0.0 | ND | ND | ND | ND | ND |
| R245C | 0.0 | 0.0 | ND | ND | ND | ND | ND |
| V246F | 3.9±0.5 | 0.2±0.1 | 1.2±0.1 | 0.011±0.004 | 27% | 50±11 | <10 |
| C290W | 0.0 | 0.0 | ND | ND | ND | ND | ND |
| C306W | 0.0 | 0.0 | ND | ND | ND | ND | ND |
| R309C | 4.9±0.3 | 0.2±0.1 | 1.0±0.1 | 0.012±0.005 | 9% | 246 ±32 | <10 |
| ^a^kinetic parameters and dissociation constants were measured as described in the methods. ^b^Metal analysis was conducted using Inductively Coupled Plasma-Mass Spectrometry (ICP-MS). Values equal to 0 means that the protein sample had Zn and Fe content equal to the blank. ND; not determined. | | | | | | | |

| **Table S11**. Mutation suppression activity^a^ of cancer-associated mutants within the [4Fe-4S] cluster of MBP-MUTYH. | | |
| --- | --- | --- |
| MUTYH | Mutation frequency (fx10^8^) | Relative change to WT |
| Empty vector (pMAL) | 5.77(4.68-6.86) | 52 |
| WT | 0.11(0.08-0.14) | 1 |
| W103C | 5.07(3.14-7.17) | 46 |
| W103R | 5.38(3.36-7.40) | 48 |
| N238S | 1.31(0.34-2.18) | 12 |
| R241Q | 2.00(1.21-2.80) | 18 |
| R241W | 4.22(2.65-5.83) | 38 |
| R245H | 2.57(1.74-3.42) | 23 |
| R245C | 3.69(2.99-4.39) | 33 |
| V246F | 0.55(0.41-0.70) | 5 |
| C290W | 6.09(2.43-9.74) | 55 |
| C306W | 5.06(3.54-6.59) | 46 |
| R309C | 1.12(0.36-1.87) | 10 |
| ^a^Mutation frequency values are shown with 95% confidence limit based on the median value. | | |

**Table S12: Reduction of restraints throughout equilibration during molecular dynamic simulations.**

| Restraints (kcal mol^-1^ Å^2^) | Time (ns) |
| --- | --- |
| 500 | 0.1 |
| 400 | 0.1 |
| 300 | 0.1 |
| 200 | 0.1 |
| 100 | 0.1 |
| 50 | 0.1 |
| 25 | 0.1 |
| 10 | 0.1 |
| 5 | 0.1 |
| 2 | 0.1 |
| 1 | 0.1 |
| 0.5 | 0.1 |
| 0 | 0.5 |

**Figure S13.-** Results of the molecular dynamic simulations of WT MUTYH, N238S and R241Q cancer-associated variants in human structure. A) Root Mean Square Deviation (RMSD), B) Energy Decomposition Analysis (EDA) with respect to residue Asn238 (in purple), with respect to residue Arg241 (in pink), C) Normal Mode Analysis (NMA) – Percentage contribution to total motion by each mode number

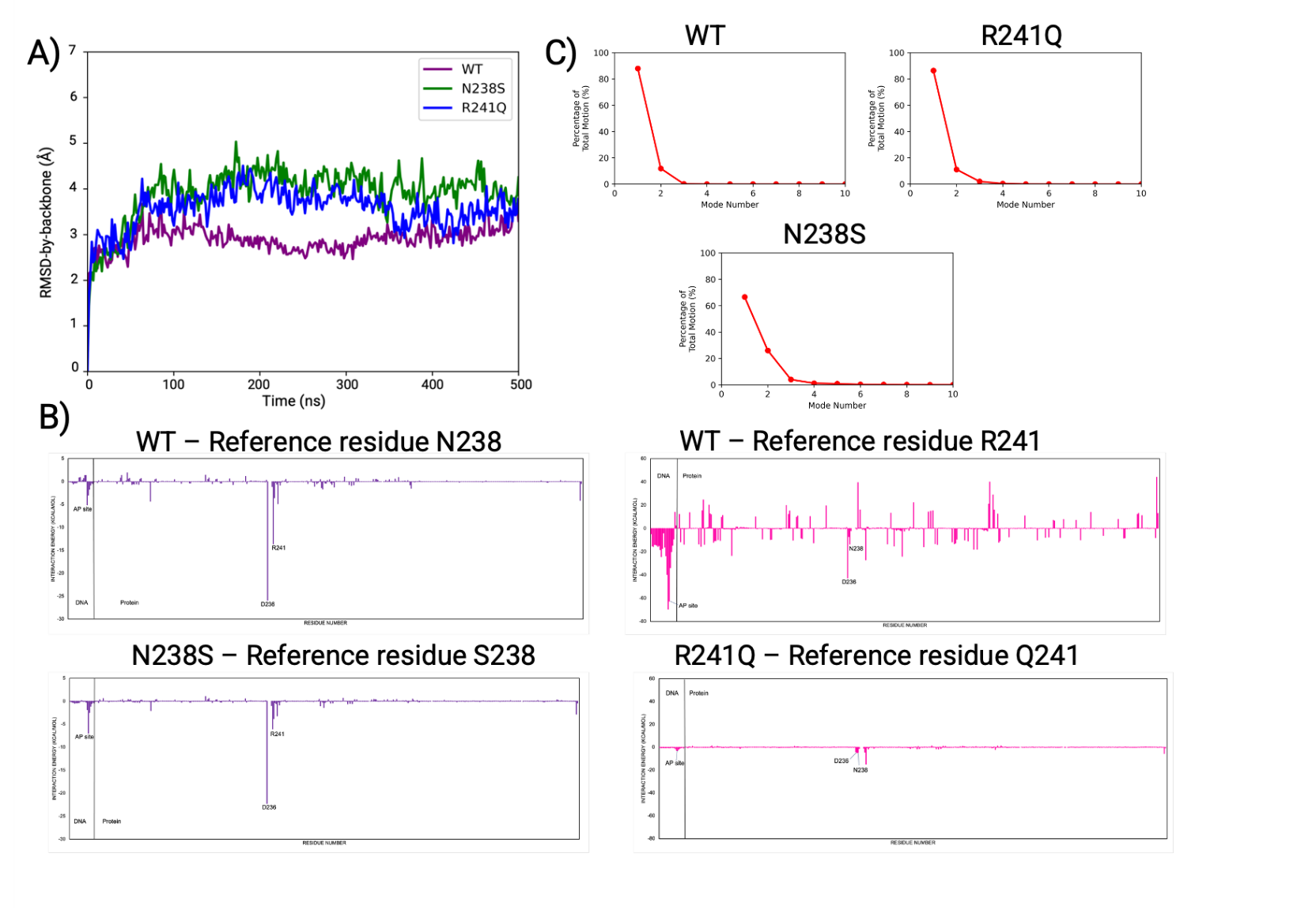

**Figure S14. -** Energy decomposition analysis of the nonbonded interactions of the MUTYH human structure with respect to the [4Fe-4S] cluster. The energy difference between (A) N238S and WT and (B) R241Q and WT are shown. Residues highlighted in red color and blue color represent the interaction energies ≥ +2 or ≤ -2 kcal mol^-1^.

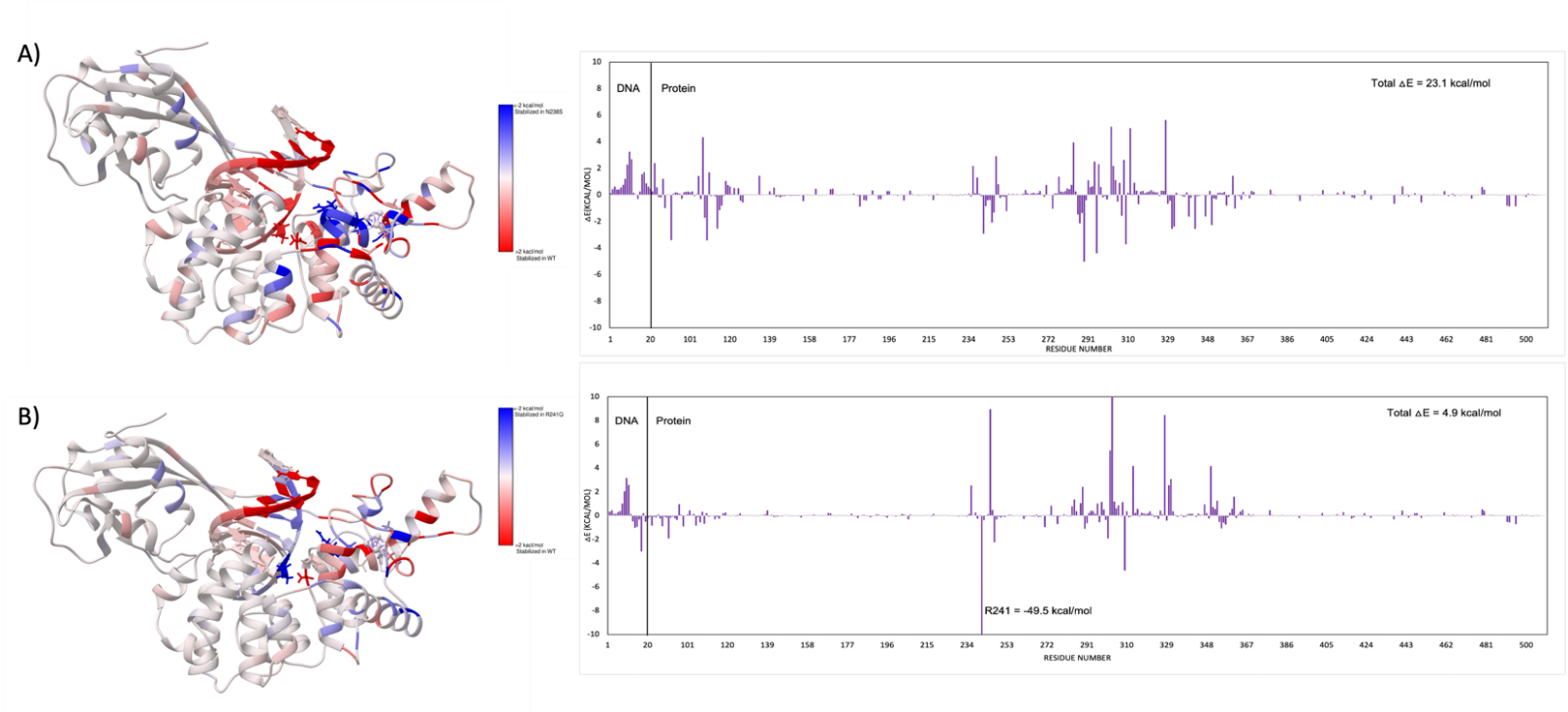

**Figure S15.-** Results of the molecular dynamic simulations of WT, N209S and R212Q cancer-associated variant in mouse Mutyh structure. The Energy Decomposition Analysis (EDA) (Left panel) shows the intermolecular non-bonded interactions (Coulomb and Vander Waals interactions; kcal/mol) between Asn209 or Arg212 (black and red values, respectively) and the rest of the residues involved. For the Network Analysis the betweenness of the nodes involved in the multi-motif bridge is shown in brown color (Right-upper panel) and the optimal path between the AP site and [4Fe-4S] cluster is displayed in green (Right-lower panel). The First mode analysis is shown on the middle where the Mutyh and DNA are illustrated in magenta and gray, respectively.
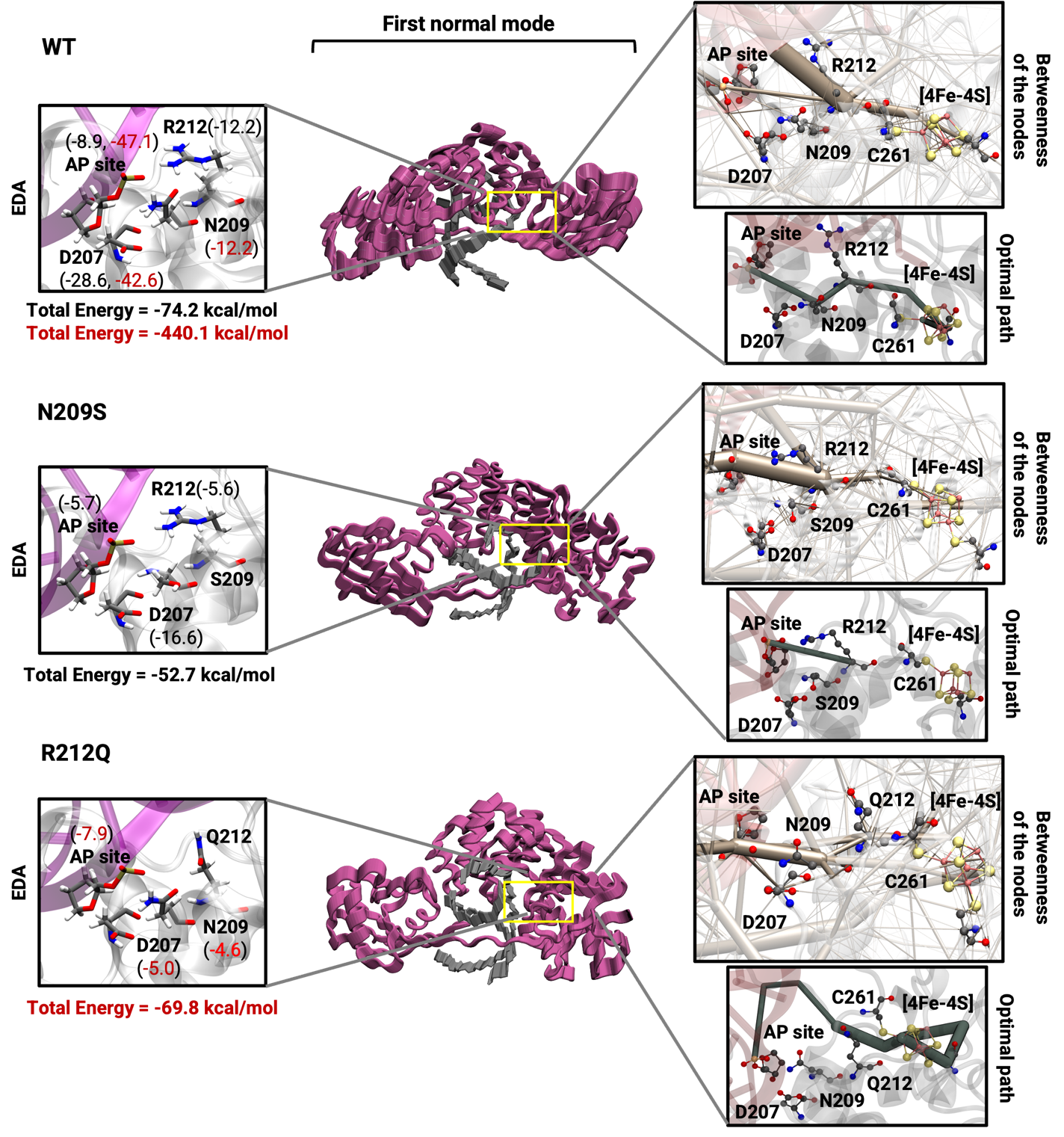

A different behavior in networks is observed in mouse Mutyh structure compared to human MUTYH structure. Figure 3 (right) shows the betweenness of nodes in brown and the optimal path between AP site and Fe-S cluster in dark green. In the WT structure, the AP site exhibits mild contacts with D207, N209 and somewhat stronger contact with R212. Conversely, the N209S mutant system shows a high betweenness, establishing a strong contact between AP site and R212. Furthermore, a strong connection is observed between AP site and N209 in the R212Q mutant structure while a connection is maintained between AP site and catalytic residue D207. Additionally, the optimal path between AP site and Fe-S cluster in the WT structure involves N209 and R212. This path is disrupted in N209S mutant structure and alternative path through the protein takes place without involving the residues in the H-bond bridge in R212Q mutant structure.

**Figure S16.-** Results of the molecular dynamic simulations of WT, N209S and R212Q cancer-associated variant in mouse Mutyh structure. A) Root Mean Square Deviation (RMSD), B) Energy Decomposition Analysis (EDA) with respect to residue Asn209 (in purple), with respect to residue Arg212 (in pink), C) Normal Mode Analysis (NMA) – Percentage contribution to total motion by each mode number.

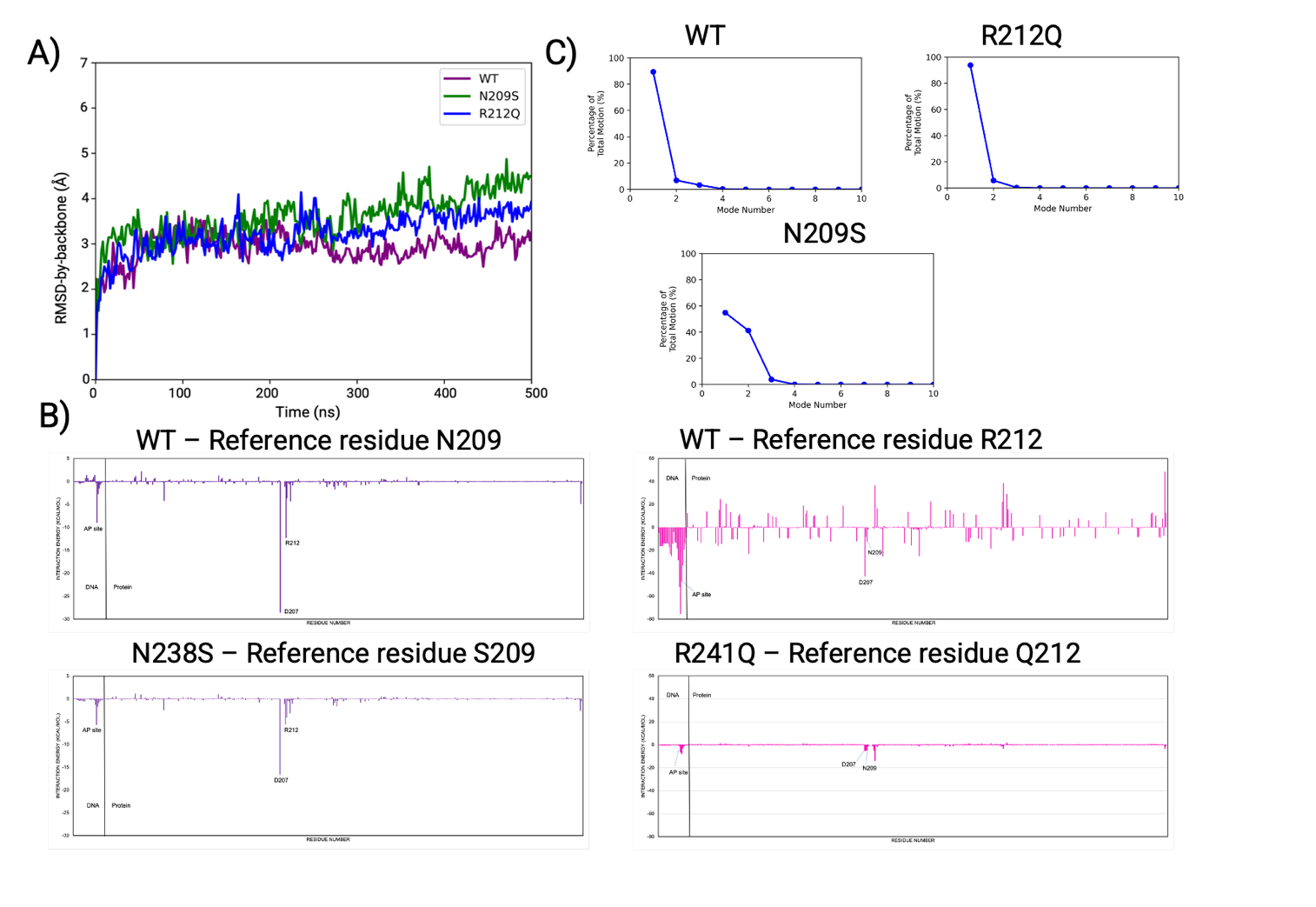

**Table S17.-** Data collection and refinement statistics for MUTYH-TSAC and R149Q GsMutY structures.

| **Data collection statistics** | | |
| --- | --- | --- |
| **PDB ID** | **8FAY** | **9BS2** |
| *Protein* | MUTYH-TSAC | R149Q GsMutY |
| *DNA* | OG:N1 | OG:THF |
| *Wavelength (Å)* | 0.97918 | 0.97741 |
| *Resolution range (Å) ** | 60.3–1.91 | 70.3–1.51 |
| *Highest resolution shell (Å)* | (2.02–1.91) | (1.55–1.51) |
| *Space group* | P2_1_2_1_2_1_ | P2_1_2_1_2_1_ |
| *Unit cell a (Å)* | 87.686 | 37.500 |
| *b (Å)* | 116.422 | 86.000 |
| *c (Å)* | 118.378 | 140.640 |
| *Total reflections*** | 484,096 (66,132) | 66,1741 (31,326) |
| *Unique reflections* | 177,017 (27,556) | 136,096 (9,284) |
| *Multiplicity* | 2.7 (2.4) | 4.8 (3.4) |
| *Completeness (%)* | 97.6 (94.1) | 98.7 (90.9) |
| *Mean I/sigma(I)* | 7.53 (0.56) | 12.2 (0.51) |
| *Wilson B-factor (Å^2^)* | 44.2 | 33.6 |
| *R-merge (%)* | 8.5 (177.9) | 5.4 (221.1) |
| *R-rim (%)^†^* | 10.4 (221.3) | 6.1 (256.7) |
| *CC1/2 (%)* | 99.8 (20.9) | 99.9 (12.4) |
| * Statistics for the highest-resolution shell are shown in parentheses.  ** Friedel mates treated as different reflections  † Redundancy independent measure of R^9^ | | |
| **Refinement statistics** | | |
| **PDB ID** | **8FAY** | **9BS2** |
| *Protein* | MUTYH-TSAC | R149Q GsMutY |
| *DNA* | OG:N1 | OG:THF |
| *Resolution range (Å) ** | 60.3–1.91 | 70.3–1.51 |
| *Highest resolution shell (Å)* | (1.98–1.91) | (1.56–1.51) |
| *Reflections*** | 174,855 (14,498) | 132,137 (12,247) |
| *R-work* | 0.180 (0.395) | 0.207 (0.424) |
| *R-free* | 0.208 (0.399) | 0.230 (0.428) |
| *Protein:DNA per asymmetric unit* | 2 | 1 |
| *TLS Groups* | N.A. | 3 |
| *Atoms, non-hydrogen* | 7492 | 3477 |
| *Protein* | 6102 | 2783 |
| *DNA* | 844 | 422 |
| *OG nucleotide* | 46 | 23 |
| *Active site nucleotide* | 22 | 11 |
| *SF4* | 16 | 16 |
| *Sulfate* | 65 | N.A. |
| *Calcium* | N.A. | 3 |
| *Solvent* | 465 | 249 |
| *RMSD bonds (Å)* | 0.012 | 0.012 |
| *RMSD angles (°)* | 1.138 | 1.179 |
| *Ramachandran favored (%)* | 97.7 | 97.4 |
| *allowed (%)* | 2.3 | 2.3 |
| *outliers (%)* | 0.0 | 0.3 |
| *Rotamer outliers (%)* | 0.5 | 0.7 |
| *MolProbity Score* | 1.02 | 1.32 |
| *Clashscore* | 2.08 | 4.24 |
| *Average B non-H (Å^2^)* | 46.9 | 41.8 |
| *Protein (Å^2^)* | 44.9 | 43.2 |
| *DNA (Å^2^)* | 58.1 | 35.2 |
| *OG nucleotide (Å^2^)* | 34.0 | 20.9 |
| *Active site nucleotide (Å^2^)* | 45.6 | 23.1 |
| *SF4 (Å^2^)* | 33.7 | 23.8 |
| *Sulfate (Å^2^)* | 68.6 | N.A. |
| *Calcium* | N.A. | 38.2 |
| *Solvent (Å^2^)* | 49.6 | 38.1 |
| ** Friedel mates treated as different reflections | | |
